# Making AI accessible for forensic DNA profile analysis

**DOI:** 10.1101/2025.06.02.656996

**Authors:** Abel KJG de Wit, Claire D Wagenaar, Nathalie AC Janssen, Brechtje Hoegen, Judith van de Wetering, Huub Hoofs, Simone Ariëns, Corina CG Benschop, Rolf JF Ypma

**Affiliations:** Division of digital and biometric traces, Netherlands Forensic Institute; Division of biological traces, Netherlands Forensic Institute

**Keywords:** artificial intelligence, deep learning, convolutional neural network, u-net, forensic DNA analysis, automated allele calling

## Abstract

Deep learning has the potential to be a powerful tool for automating allele calling in forensic DNA analysis. Studies to date have relied on bespoke model architecture and painstaking manual annotations to train models, which makes it challenging for other researchers to work with these techniques.

In this study, we explore the possibility of training a well-performing model using data gathered as part of casework, and employing a widely adopted architecture: the U-Net. In this approach, annotations are created from alleles called during casework. The model, dubbed ‘DNANet’, then classifies each scan point in the electropherogram (EPG) as part of an allele or non-allele, building on the task of segmentation in computer vision. We evaluate performance on unseen case data and on independent mixture research data, taking analyst annotations as ground-truth. We further compare DNANet’s performance with analyst performance on the research data, taking actual donor alleles as ground-truth.

DNANet reached an F1 score of 0.971 on analyst annotated alleles on case data not seen during training, and 0.982 on the research data. On actual donor alleles, DNANet reached an F1 score of 0.962, equal to the F1 score computed from analyst annotations.

Our results show that DNANet’s performance is comparable to human annotations following standard procedures. This illustrates the potential for obtaining good results with standard data and architecture. Future work may focus on what aspects of data, annotations or model architecture are key in shaping performance. We make our code, model weights and research data publicly available to aid the community. Lastly, we call for an effort to establish a standardized benchmark to aid in quantitative comparisons between allele calling systems.

## Introduction

Machine learning methods, in particular deep learning methods, form a promising technique for the task of allele calling in forensic DNA analysis (1–10). Traditional methods for analyzing electrophoresis data often rely on manual inspection and rule-based systems, which can be time-consuming, open to human error and can lead to inconsistencies between analysts. In contrast, deep learning models can learn complex patterns directly from raw data, enabling faster and more consistent results.

Despite the promise of applying deep learning to allele recognition in forensic DNA analysis, papers have remained limited to the pioneering work of a single group. Taylor et al. introduced a system of ten separate neural networks, each dedicated to a specific dye lane or locus, to classify electropherogram signals into categories such as alleles, stutter, pull-up, and baseline noise (2, 3). This modular design was necessary to account for locus-specific signal behaviors, but it also fragmented the learning process and required redundant training across networks. Later improvements led to a unified multi-head convolutional neural network that could share early feature representations across dye lanes while branching into specialized outputs (1). This model worked exceptionally well, whilst forgoing the conventional approach of a hierarchy of filters learning consecutively higher-order structures for a single filter with a width of 200 scan points. All these models relied on painstaking manual annotations of hundreds of profiles, specifying scan points to belong to one of eight categories (baseline, allele, pull-up, and five stutter types). The combination of handcrafted pipelines and manual annotation make the models hard to build on or reproduce, especially as data are often too sensitive to share in this field.

In this work, we investigate two simplifications. First, we use the U-Net architecture as the basis of our model (11). This architecture is ubiquitous in the task of (bio-medical) image segmentation (12, 13). Segmentation involves assigning a category label to each part of a continuous input—such as labeling each pixel in an image or each data point in a signal. In the context of DNA profiling, segmentation would be classifying individual EPG scan points as being part of an allele. Second, we rely on data gathered as part of casework, namely the called alleles. Thus, we avoid the intermediate step of classifying various forms of stutter and pull-ups employed before, allowing for a much lower entrance barrier to developing deep learning methods. However, this also means there is less information per profile for the model to learn from.

This study describes the development and performance evaluation of our deep learning model for DNA profile analysis, dubbed ‘DNANet’, and discusses directions for future research. The code, model weights and research data are made publicly available to aid laboratories seeking to develop or evaluate deep learning for forensic DNA analysis.

## Background

### Segmentation

A universal task in computer vision is to automatically determine exactly where in an image an object of interest is located. This task is referred to as segmentation. Applications of segmentation range from the localization of tumour cells in medical images to the detection of objects by autonomous vehicles. An example application in the forensic field is to automatically find traces of interest, such as fibres in microscopy images (see Fig. 1) (14).

**Fig. 1.**
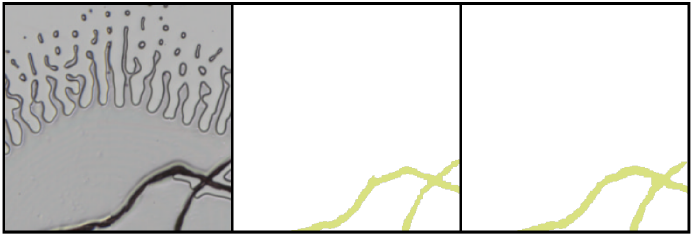
Example of a segmentation task in the forensic field. (Left) Microscopy image with fibre of interest, (middle) manual annotation of the fibre and (right) pixels indicated by the algorithm to compose the fibre. Taken with permission from Figure 2f in (14).

Calling alleles from an electropherogram can be seen as a segmentation task. Natural images, e.g. photographs, consist of a matrix of pixels, and the algorithm classifies each of these pixels as being or not being part of an object of interest. The electropherogram consists of measurements in Relative Fluorescence Units (RFU) across five dyes (the sixth is an internal standard) and can thus be seen as an ‘image’ of five by several thousand scan points. Allele calling is then akin to classifying each of these scan points (‘pixels’) as being part or not being part of an actual allele. It thus seems a natural approach to apply algorithms well suited to image segmentation to allele calling.

### U-Net Architecture

One of the most commonly used architectures for segmentation tasks in machine learning is the UNet (11, 12). Originally developed for medical image analysis, a U-Net is well suited to tasks where each part of a signal or image needs to be labeled individually. This is especially relevant for data like electropherograms, where individual alleles need to be detected. The U-Net architecture follows a characteristic U-shape with two main components: an encoder and a decoder (see Fig. 2). The encoder progressively reduces the resolution of the input, summarizing broader patterns and compressing the data into abstract, informative features. This process is similar to how a human might first get a general sense of a DNA profile before zooming in on specific peaks. However, in doing so, fine details can be lost. To recover those details, the decoder upsamples the compressed data back to its original resolution. At each step, the decoder receives skip connections from the encoder—direct links that reintroduce earlier, higher-resolution information. This design allows the network to combine the broader context with precise local structure, resulting in more accurate and finegrained segmentations.

**Fig. 2.**
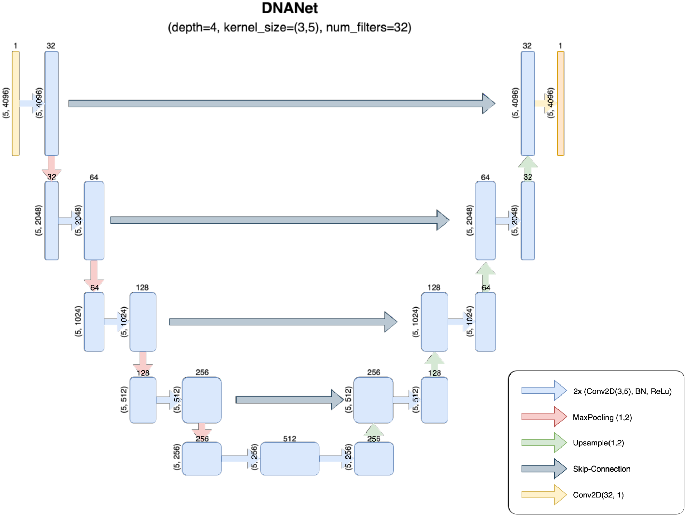
Schematic drawing of DNANet’s architecture. The downward stream of blocks on the left form the encoder, the upward stream of blocks on the right the decoder. Together this forms the distinctive ‘U’ shape of a U-Net.

## Materials & Methods

### Data

The dataset used in this study consists of two sources: casework and research. All are PowerPlex® Fusion 6C (PPF6C) profiles. The casework data are used for training and evaluating DNANet. The research data are used as an independent dataset for additional testing, and are published online. The casework data reflect the diversity and complexity encountered in real forensic practice, including variation in number of contributors, mixture proportions, degradation, artifacts, etc. The casework data were analysed by DNA analysts during the case following standard laboratory protocols. In total, the casework dataset comprised 70722 profiles.

The research profiles have been previously described (15). In short, they comprise 120 mixtures from 30 different donors, spanning different mixture proportions and 2 to 5 donors per mixture. Three replicates were measured per mixture. Ten profiles did not pass quality control, resulting in a dataset of 350 profiles.

For each profile, we have a) an .hid file containing raw fluorescence signal values (RFU) measured, b) the ladder associated with the profile and c) the set of alleles called by the analyst in the case. For the research profiles, we additionally know the genotypes for each of the donors to the sample. Manual DNA profile analysis was performed in GeneMarker HID v2.9.8, using dye specific analytical thresholds, stutter filters and a minimum heterozygote peak imbalance percentage. Analysts check the peak shape and remove the peak or call it as an allele, possibly renaming it.

### Data preprocessing

We use the ladders and internal size standard to size each of the profiles, using a piecewise linear transformation. We then represent each profile as a 5 × 4096 matrix, where the five rows correspond to the RFU from the remaining dye channels. We create the scan points by resampling between base-pair position 65 and 475, which correspond to the first and last peak of the internal standard, cutting out any primer flare. We specifically use 4096 data points, rather than a previously used 5000 (1), for the simple reason that having a power of two makes it easier to apply the U-Net. No additional preprocessing, normalization, or augmentation is applied.

Using the set of called alleles, we construct for each profile a corresponding 5 × 4096 matrix of annotations, indicating for each scan point whether it is part of an allele (1) or not (0). These binary labels are used to supervise the training of DNANet. In our first attempt to do this, we simply took the corresponding bin for each called allele, appropriately sized to arrive at the relevant scan points. In practice, the actual peak shape does not correspond precisely to the scan points that make up the bin (see Fig. 3A). This makes it harder for the model to learn that it should be looking at the peak. We therefore explored two heuristics to improve the labels. To do this, we wrote a simple algorithm to find the peak that falls (partially) within this bin. We then labeled either the scan point at the very top of this peak (Fig. 3B) or all scan points the peak consists of, by finding the first local minimum on either side of the peak (Fig. 3C). We found that the first of these two heuristics led to highest performance, and use this approach for generating annotations from called alleles.

**Fig. 3.**
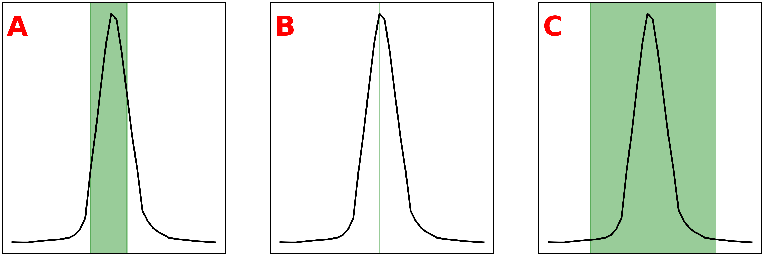
Different annotations (green) generated for the same example called allele (black line). (A) Scan points falling in the bin associated with the allele are used as annotation. Shown is an example where bin and peak don’t neatly line up, which regularly happens. (B) The scan point with maximal RFU in the peak is used as annotation. (C) All scan points in the peak are used as annotation. These are determined by finding the first local minimum on either side of the peak.

### Model

For this study, we implemented the well-known UNet architecture. U-Net is a flexible architecture with several tunable parameters that influence its capacity to model spatial patterns and its sensitivity to features of varying scale. In our implementation, we explored three key architectural parameters that govern the structure and performance of the network:

- **Network Depth**: This parameter determines the number of downsampling and upsampling levels in the UNet. Increasing the depth allows the network to capture broader contextual information, as the receptive field expands with each level. However, deeper networks also require more computational resources and may risk overfitting if training data is limited.
- **Kernel Size**: The size of the convolutional kernels affects how local features are aggregated. Larger kernels can capture wider patterns in a single layer, which may be beneficial for identifying broader peak shapes or signal artifacts. Smaller kernels, on the other hand, emphasize fine-grained detail and tend to generalize better in noisy environments.
- **Number of Filters**: This parameter controls the number of feature maps learned at each level of the network. Higher numbers increase the representational capacity of the model, allowing it to learn more nuanced signal characteristics. However, this also increases the number of parameters and computational cost.
- **Model Training**. To identify suitable hyperparameters, we conducted a grid search over key architectural and training parameters, optimizing the scan point level F1-score (see ‘Evaluation’). The following hyperparameters were varied:
- **Network depth** (i.e., number of downsampling/upsampling steps): *{*2, 3, 4*}*
- **Kernel size**: *{*(1, 3), (3, 5)*}*
- **Number of filters** in the initial convolutional layer: *{*4, 8, 16, 32, 64*}*

In total, this resulted in 90 unique model configurations. The casework dataset was split into 80% for training and 20% for testing, stratified by robot (four ABI3500xLs are employed in casework). Then the 80% for training was split again into 80/20 for training and grid search evaluation. Each model was trained and evaluated on the same data split to ensure comparability. This double splitting ensured the 20% test data were never used in the grid search. Optimization was performed using the Adam optimizer (16) with a starting learning rate of 0.0001, combined with an exponential learning rate scheduler (`torch.optim.lr_scheduler.ExponentialLR`) using a decay factor (gamma) of 0.8. This gradually reduced the learning rate over epochs to stabilize training as convergence was approached. We optimized on standard Dice loss, an often used loss function for segmentation.

### Allele calling

The output of DNANet is a 5 × 4096 matrix of predictions—one prediction for each scan point (see Fig. 4 for a visualisation). In the final step, these predictions are converted back into called alleles based on the locations of predicted peaks. To do this, we apply the following heuristic. The heuristic processes each dye-lane individually. It identifies all scan points with a prediction score greater than or equal to 0.5. These are then naively grouped based on contiguity—each time a prediction drops below the threshold, a new group is started. For each group, each scan point is translated following the internal standard. From that translation, the group’s mean is computed, referred to as the “mean base-pair.” Using the ladder and internal standard per profile that maps alleles to expected base-pair (bp) positions (e.g., AMEL X = 18.5 bp), each group’s mean base-pair is matched to the allele with the nearest base-pair position. The final result is a list of alleles that DNANet predicts to be present in the DNA profile.

**Fig. 4.**
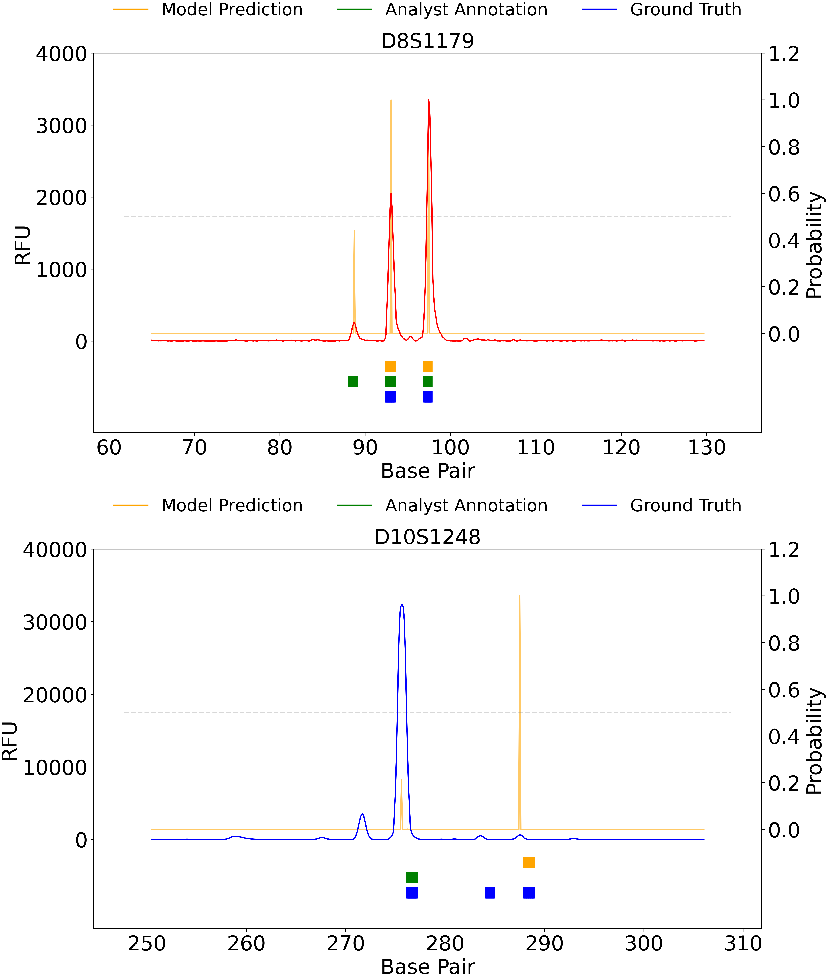
Examples of (orange; right axis) scan point level predictions generated by DNANet from the (red; left axis) RFU signal. The threshold of 0.5 for calling alleles is visualized as a dotted horizontal line. The coloured bars underneath the line plot show (blue) the donor alleles for this locus, (green) the alleles called by the analyst and (orange) the alleles called by DNANet. For visibility, RFU=0 is plotted slightly lower than probability=0. (top) Predictions for locus D8S1179 from research profile 3D2. The prediction reaches 1 for the two donor alleles, is around 0.4 for the stutter (at around x=230) and is 0 otherwise. This results in DNANet calling the two alleles and nearly (incorrectly) calling the stutter. Here, the analyst did call the stutter. (bottom) Predictions for locus SE33 from profile 1E3. Here, one allele (bp 372; on stutter position) was missed by both DNANet and the analyst. The analyst missed an allele at the start of the locus (bp 317). Plot can be reproduced using the `create_paper_figures.py` script available online.

Note that this heuristic can be further optimized. For example, predicting a scan point outside of a marker region as allelic will incorrectly lead to calling a peak as an allele. Furthermore, two alleles whose peak shapes overlap may well be called as a single allele using this approach. Also note that there is an interaction with the method selected to create annotations from the called alleles (see Fig 3), our current method of only assigning a positive value to the top scan point may result in less sensitivity to the heuristic used here. We have not explored these matters at great depth as the focus of the current paper is on the deep learning technique. The allele calling serves as a way to measure, albeit crudely, its performance at allele calling.

### Evaluation

Model performance was assessed at three levels: for individual scan points, individual alleles, and for individual donors. The first reflects how accurately DNANet labels individual scan points, the second shows whether alleles were correctly called. For the third, we compute LRs for actual donors using the called alleles. This multi-layered evaluation reflects both the fine-grained accuracy of the segmentation task and the practical relevance of correctly identifying allelic peaks. We compute the following standard measures:

- **Precision**: the proportion of predicted alleles that correctly match a true allele.
- **Recall**: the proportion of true alleles that were correctly identified by DNANet.
- **F1-score**: the harmonic mean of allele-level precision and recall.

At the scan point level we report only the F1-score, at the allele level we report all three. We report these metrics both for the test case data and for the independent research data. For the research data, the ground-truth is available in the form of the alleles of the actual donors to the samples. This allows us to compare DNANet’s and analyst’s ability to call alleles, something not possible for the case data. We compute the above allele-level metrics for both DNANet and analyst called alleles, taking all donor alleles as ground-truth. Note that 100% accuracy will likely not be possible for this dataset, as not all alleles are visible in the electropherogram (drop-out).

Finally, we compute the likelihood ratios (LRs) for all donors to the research data for 2p, 3p and 4p mixtures, both using the alleles called by the analyst and those called by DNANet (n=787 LRs). LRs were computed using DNAStatistX v2.3.5, using the known NOC under the propositions. The stutter model was off and the degradation model was on.

### Open science

The case data cannot be shared because of sensitive information. However, to be transparent and help others set up these kind of analyses, we provide at github.com/NetherlandsForensicInstitute/dnanet:

- The research data in raw and analysed form,
- The model weights, trained on case data,
- The Python source code for reading the data, and training and evaluating models.

This should lower the barrier for others to run these analyses. The code has been set up to make it relatively straightforward to train on one’s own data.

## Results

To better understand what DNANet outputs, Fig. 4 visualizes the output for two loci from two different profiles from the research data (see Fig. S1 and S2 for full output). It can be seen how DNANet computes a number between 0 and 1 for each scan point, indicating whether this scan point is the top of an allele. In our approach, predictions higher than 0.5 will lead to an allele being called. We found that most often DNANet outputs a score close to 1 for alleles and close to 0 otherwise. Exceptions are more ‘difficult’ parts of the electropherogram, such as the relatively high stutter in Fig. 4, which can lead to intermediate scores. The figure also shows the alleles called by the analyst, and the donor alleles. Both analyst and DNANet often, but not always, call the correct alleles. The full visualisation for these two profiles corresponding to Fig.4 are given in supplementary figures S1 and S2. Figures can be reproduced and created for other profiles by running `create_paper_figures.py` from the github repository.

From our grid search, the following combination of parameters provided best results: 32 filters, a network depth of 4, kernel size of (3, 5) and 3 training epochs. The resulting model achieved an F1-score in classifying individual scan points of 0.966. Using these classifications to call alleles led to a recall of 0.960, precision of 0.983 and thus F1-score of 0.971 on the casework data test dataset (Table 1).

**Table 1.** DNANet performance on casework and research data, taking analyst annotations as ground-truth.

|  | Casework data | Research data |
| --- | --- | --- |
| Recall | 0.960 | 0.980 |
| Precision | 0.983 | 0.984 |
| F1 | 0.971 | 0.982 |
| Scan point F1 | 0.966 | 0.970 |

Table 2 shows the results when evaluating both DNANet and analyst annotations against ground-truth of donor alleles for the research data. We find that performance is very similar, with the F1-score for both at 0.962. The majority of mismatches between allele calls and ground-truth were alleles with low RFU, often at stutter position. We found one exception, where DNANet fails to call an allele with high RFU, which we visualize in Fig. 5. It is unclear why the model misses this allele.

**Table 2.** DNANet and analyst performance on research data, taking actual donor alleles as ground-truth.

|  | Analyst | DNANet |
| --- | --- | --- |
| Recall | 0.936 | 0.934 |
| Precision | 0.990 | 0.992 |
| F1 | 0.962 | 0.962 |

**Fig. 5.**
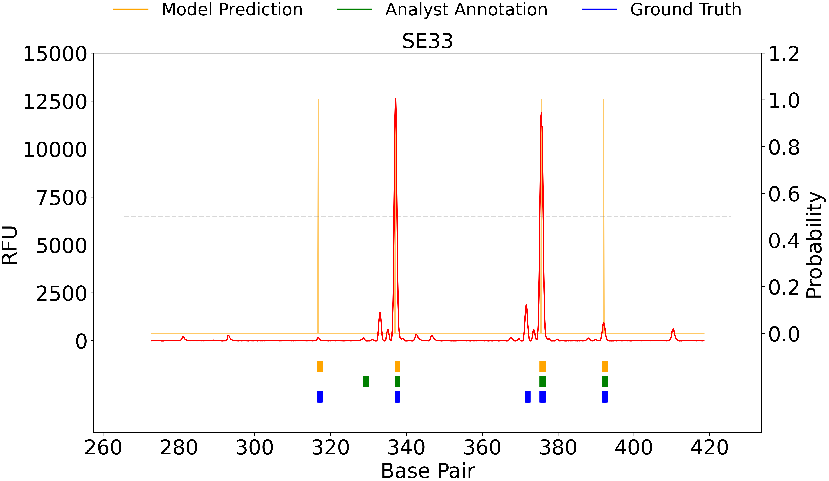
Visualisation of predictions generated for marker D10S1248 of profile 1E3, selected as DNANet missed an allele with high RFU. Shown are (blue line) the RFU, (yellow line) DNANet prediction per scan point, (green square) analyst called peaks (yellow square) DNANet called alleles and (blue square) ground-truth alleles. There are three alleles present, where the first has very high RFU (>30000) and the latter two are low. The analyst only called the high peak. Surprisingly, DNANet only called the last peak. Its prediction for the high peak was around 0.2, not clearing the threshold of 0.5.

Finally, we examined DNANet’s performance by comparing LRs obtained for H1-true donors from DNA profiles, analysed either by DNANet or an analyst. Figure 6 shows the LRs found for the actual donors to the research DNA profiles. The figure shows that, despite the differences in allele calling, the resulting LRs are very similar. For instance, for 78% of H1-true donors the LRs based on analyst and DNANet annotations differ by less than a factor of 10.

**Fig. 6.**
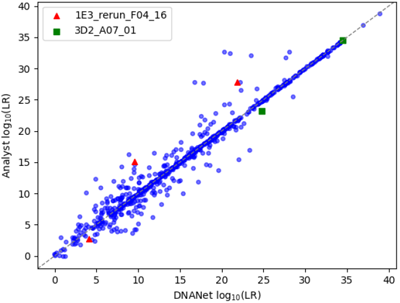
Likelihood ratios found for actual donors to the two-person, three-person and four-person mixture research data, for Analyst or DNANet called alleles. The three H1-true LRs for profiles 1E3 (Fig. 4 bottom; Fig. 5) are shown as red triangles, the two H1-true alleles for profile 3D2 (Fig. 4 top) as green squares.

## Discussion

In this manuscript, we showed how a well-performing AI model for allele calling can be trained using a standard model architecture, and data and annotations obtained during standard casework. The model, ‘DNANet’, is able to learn from analyst annotation patterns and shows comparable performance as these analysts. We have made the model weights, source code and research data publicly available.

Although we have shown a straightforward U-Net performs well on forensic DNA analysis, more suitable architectures may exist. Here, we cast the problem of allele calling as a segmentation task, for which the architecture is well suited. However, some crucial differences exist between allele calling and segmentation. First, there is no ‘logical’ ordering of the six dyes in electropherograms, and after sizing the only relevant relation between the dyes are artefacts such as pull-ups. Thus, it may be beneficial to have distinct filters for the vertical and horizontal direction, sharing knowledge on the structure of pull-ups across dyes. Second, in natural images objects of interest can be very small or large, e.g. a dog in the foreground or background. In other words, global features are relevant for evaluating local structures. In electropherograms objects are always at the same scale. Thus, the U-Net’s strength of representing high-level features in its bottleneck may be unnecessary or even harmful for DNA profiles. It may be more logical to keep the dye dimension intact throughout the model such as done by Taylor (1). Third, as all alleles are simple short horizontal lines in the 5 × 4096 matrix, additional inspiration for solutions may be found in the domain of object recognition (17).

This study focused on investigating to what extent casework data and standard architecture were suitable for deep learning model development. There are several reasons why the DNANet is not ready for real-life application. First, an algorithm that occasionally misses high peaks such as shown in Fig. 5 without a clear reason is not ready for casework. Second, DNANet may perform poorly on rare technical artifacts. For example, we noticed the model performed poorly on profiles with strong baseline drift. This is not something we adjust for in preprocessing and happens relatively rarely - the model has not properly learned to handle such a profile and assigned positive allele labels to nearly all scan points. Third, in practice it is often the smaller peaks that are of most interest in allele calling, as the high peaks are relatively straightforward to call. As our laboratory procedures use analytical thresholds, DNANet has learned that peaks below this threshold are not alleles. This mimics analyst behaviour well but prohibits us from increasing the model’s sensitivity to minor donors. Both of the above issues may be handled in future work by oversampling relevant profiles, respectively those with baseline drift and analysed at lower or no threshold. Fourth, more broadly, we have no ground-truth data for our casework profiles. This means it will be hard for DNANet to outperform human analysts, and impossible for us to measure this if it happens. We have seen some indications that DNANet may find human errors (Fig. 4), as was found by Taylor (1).

A promising complementary approach to training the model would be to use synthetic data. Taylor and Humphries showed how a generative model can be constructed that generates lifelike electropherograms with a pre-specified number of donors and mixture contributions (18). Such a model allows for training on a large collection of non-sensitive profiles that feature precise ‘ground-truth’ labels and are specifically tailored to be ‘difficult’, e.g. having many small peaks. Of course, validation on real lab-specific data will still be needed as there may be subtle differences between generated and real data. It is quite possible that training on a combination of synthetic, experimental and real data will yield the best model.

Although our results are broadly comparable to previous work, an in-depth comparison is currently not possible. This is because other datasets and other evaluation metrics are used across studies. For example, Taylor (1) annotates all scan points that make up a peak and we only annotate the top, therefore scan point F1 score of 0.868 cannot be directly compared to our F1 score of 0.966. Most of their errors seem to be made in the tails of the peaks, which is not an issue with our annotation that only labels the top of the peak. Their allele F1 score of 0.995 (computed from their mention of 16 false positives on 1700 alleles) is more amenable to comparison, and clearly higher than our 0.971. This may well be due to their model recognising low alleles better, given the nature of the training data. Likewise, comparison to Tan et al. (10) is not straightforward. They take the identification of peaks as a given and then focus on the simpler task of classifying the peak, which makes direct comparison difficult. Moreover, all these studies are computed on different datasets.

Standardized benchmarks would be the best way to evaluate and compare different AI models and systems for allele calling. Unlike other machine learning domains where large, labeled datasets are available for validation and comparison, forensic DNA analysis lacks such resources due to the highly sensitive and individualized nature of the data and the relative novelty of these data-driven approaches. Such a benchmark may perhaps be created from the PROVEDIt dataset, to our knowledge the only freely available comprehensive dataset with ground-truth labels (19). Future work may thus focus on providing code to evaluate and report model performance on these annotated profiles in a systematic way. Note that this would also require evaluating model performance across kits, which is an interesting venue of future work.

Providing an explanation of the AI model’s decision-making process is an important step toward building trust in its use, especially in a forensic context (20). For example, recent work by Elborough et al. showed techniques from explainable AI can be used to interrogate what features of a profile an AI model uses to classify a certain peak as allelic or pullup (21). Tan et al. take a different approach, training simpler machine learning algorithms to perform the same task (10). While it is important to acknowledge that any explanation of an AI model will be inherently limited—due to the complexity of the underlying algorithms—offering insights into how the model reaches its conclusions can help mitigate concerns, foster transparency and help construct better models.

Lastly, a trained deep learning model can serve as a foundation for related tasks in forensic DNA analysis. This is similar to how models like chatGPT are generic text models that can be used for many different tasks. By learning generalizable features from electrophoresis data the model acquires an internal representation that can be fine-tuned for downstream objectives, such as number of contributor estimation (22). This transferability reduces the need for large, task-specific datasets, as such a model only has to be trained once. This would make AI models not only a targeted solution but also a stepping stone toward a broader suite of intelligent forensic analysis methods.

## Conclusion

In this manuscript, we showed how a well-performing AI model for allele calling can be trained using a standard model architecture and data and annotations obtained during casework. We hope this finding, coupled with the publicly available code, model weights and example research data, will lower the bar for others to pursue AI applications.

## ACKNOWLEDGEMENTS

We thank Duncan Taylor for an insightful discussion and useful advice, Yann Chovory for sharing data extraction scripts, Edwin Rijgersberg and Leandra Swiers for help with models and data formats and Francisca Duijs for help in technical interpretation of profiles.

## Supplementary Information: Full profile plots

Plots for all profiles can be created using create_paper_figures.py.

**Supplementary Figure S1.**
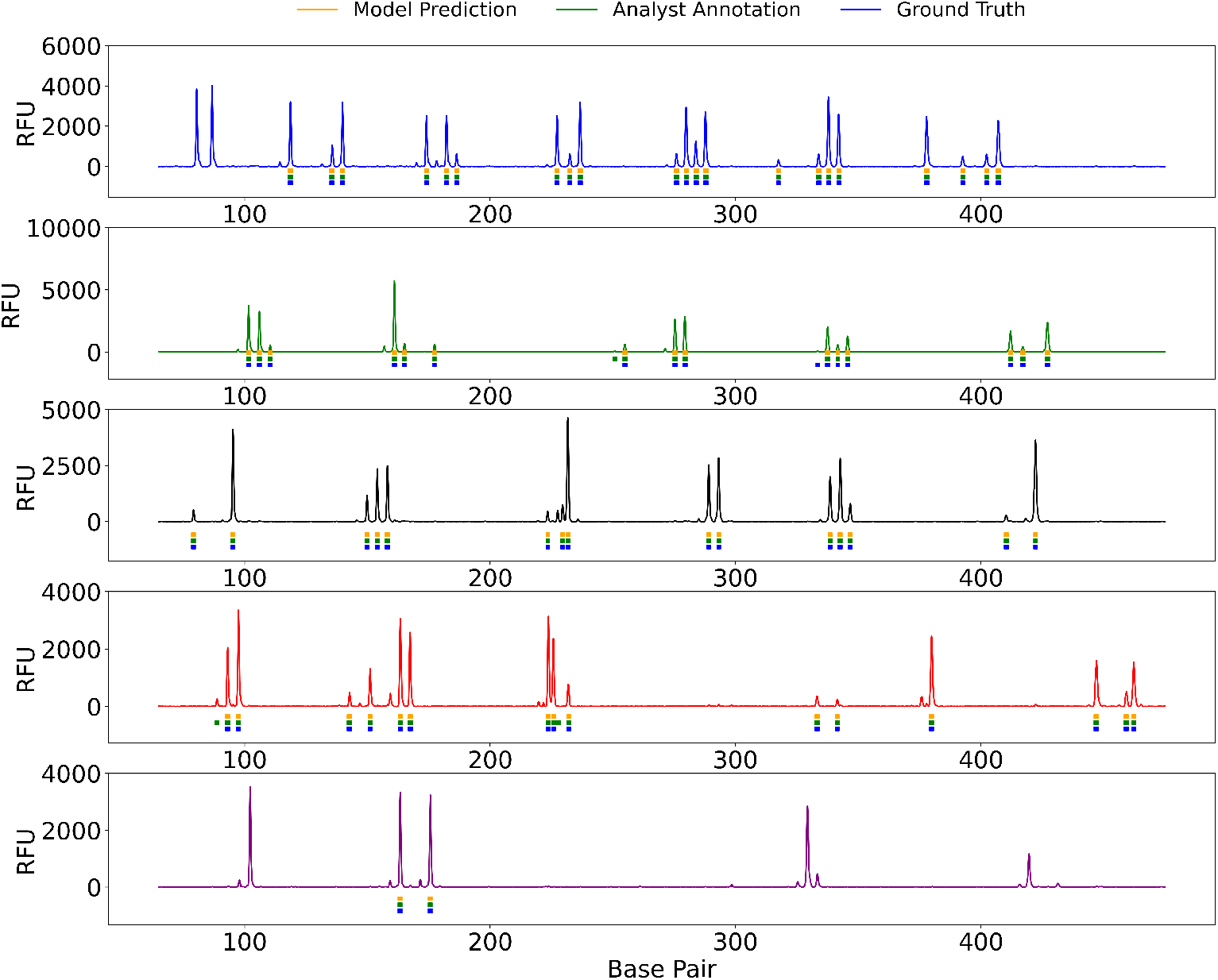
The electropherogram for sample 3D2 from the research data, corresponding to Fig. 4 from the main text. Underneath the RFU are shown (orange) the alleles called by the model, (green) alleles called by the analyst and (blue) the donor alleles. Note that ground-truth alleles were only available for the autosomal alleles, leading to missing blue bars for AMEL (top left marker) and the y-chromosomal markers in the bottom dye. In this profile, all donor alleles are called by both model and analyst, except for one allele at stutter position in CSF1PO (green dye), that was missed by both. Additionally, the analyst incorrectly called an allele at D8S1179 (also shown in Fig. 4 main text).

**Supplementary Figure S2.**
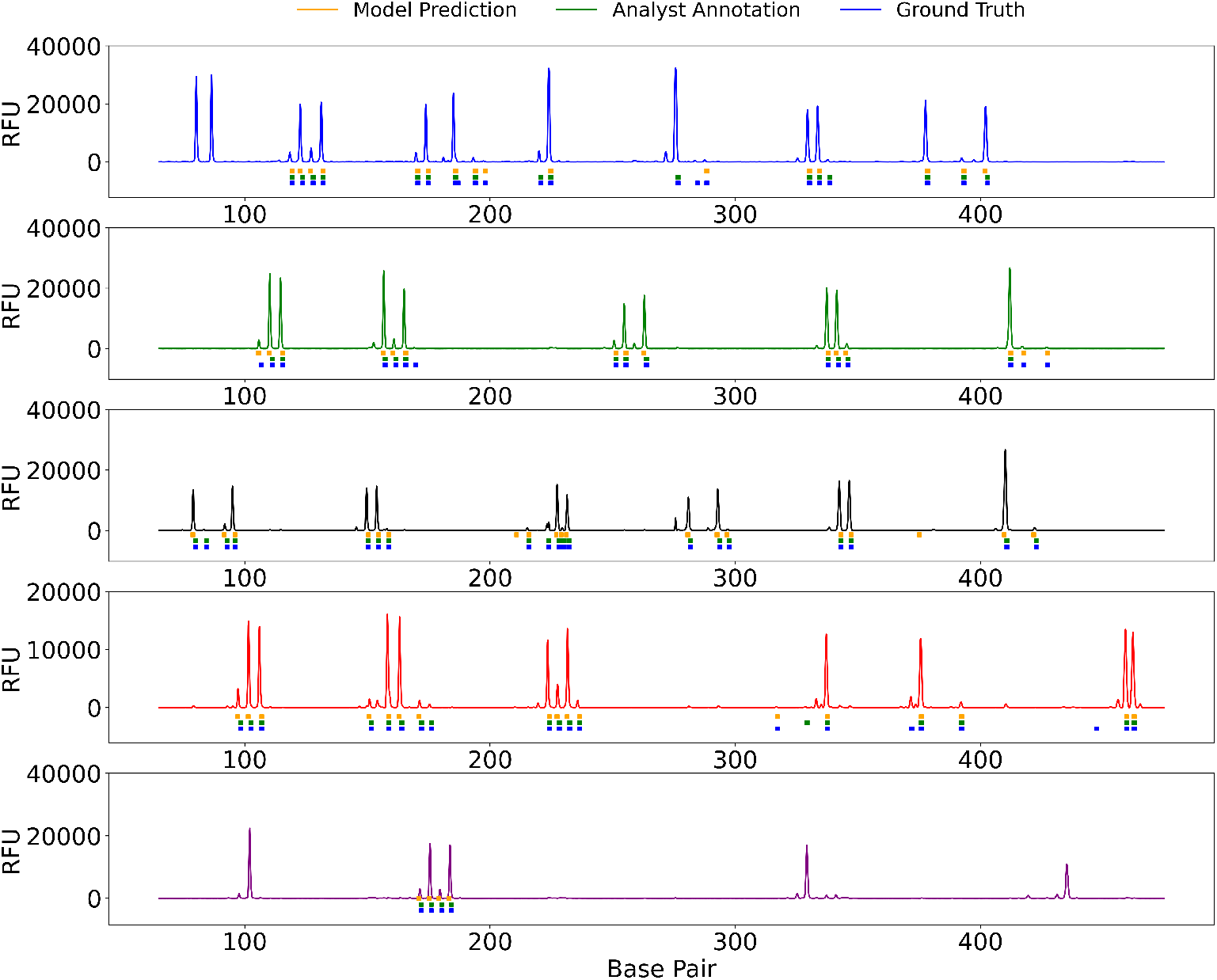
The electropherogram for sample 1E3 from the research data, corresponding to Fig 4 bottom and Fig. 5 from the main text. Underneath the RFU are shown (orange) the alleles called by the model, (green) alleles called by the analyst and (blue) the donor alleles. Note that ground-truth alleles were only available for the autosomal alleles, leading to missing blue bars for AMEL (top left marker) and the y-chromosomal markers in the bottom dye. In this profile, most, but not all, donor alleles are called by both model and analyst. The F1-score for DNANet for this profile is 0.90, the lowest score seen across profiles.

## Notes

https://github.com/NetherlandsForensicInstitute/DNANet

### Competing Interest Statement

The authors have declared no competing interest.

